# The Hidden Model Space of Phylogenetic State-Dependent Diversification Models (SSEs): Congruence, Challenges, and Opportunities

**DOI:** 10.1101/2022.07.04.498736

**Authors:** Sergei Tarasov, Josef Uyeda

**Author notes:** The authors declare no competing interests.

## Abstract

A recent study (Louca and Pennell, 2020) spotlighted the issue of model congruence, or asymptotic unidentifiability, in time-dependent birth-death models used for reconstructing species diversification histories on phylogenetic trees. The present work investigates this issue in state-dependent speciation and extinction (SSE) models, commonly used to study trait-dependent diversification. We found that model unidentifiability is universal due to hidden states, with every SSE belonging to an infinite congruence class. Notably, any trait-independent model is congruent with trait-dependent models, raising concerns for hypothesis testing. To address this, we propose an analytical solution that resolves model selection within a congruence class. Our findings show that this type of congruence is the only one possible, and with our solution in place, model unidentifiability in SSEs becomes absolutely harmless for inference. However, model selection across congruence classes remains challenging due to extremely high false positive rates. The discovered congruence offers a clear explanation of this issue and suggests potential ways forward.

---

The dramatic variation in the species richness of clades across the tree of life has lead to widespread interest in the potential drivers of diversification. State-dependent speciation and extinction (SSE) models have emerged as powerful tools for testing these hypotheses, allowing researchers to explicitly model relationships between traits and diversification rates on phylogenies (1, 2).

However, the underlying complexity of the diversification process poses a challenge when drawing reliable inferences from SSE models. Any reasonably-sized clade is likely to have factors influencing diversification that are not completely captured by the model or the focal trait. These hidden factors have been shown to mislead inference using SSEs (3, 4). To address this challenge, SSEs have adopted hidden states to model the heterogeneity of diversification processes, which may or may not be associated with the focal trait (4–6). While such SSEs advance our ability to reconstruct biologically-realistic diversification, the limitations regarding the use of hidden states and their optimal number remain uncertain.

Recently, Louca and Pennell [LP] (7) demonstrated the limitations in reconstructing diversification using time-dependent birth-death models, which are similar to SSEs but do not include a trait-dependent variation in rates (1). These limitations stem from model asymptotic unidentifiability (8) or congruence. Specifically, LP showed that any given timetree could be explained by an infinite number of alternative but congruent diversification scenarios, all yielding identical likelihood scores, regardless of the amount of observed data. The model congruence persists under any sampling fraction of species (7), even it varies over time (9), thereby limiting our ability to accurately reconstruct the true diversification history.

This discovery has sparked considerable interest in the phylogenetic community, leading to novel methods for addressing the problem. Some of these methods proposes a thorough exploration of congruent diversification scenarios (10–12), while others recommend imposing mild restrictions on extinction and speciation functions (13, 14). Additionally, an extended version of this time-dependent process, the fossilized birth-death model, has been found to be identifiable (15).

While LP’s study focused on time-dependent models, they proposed that similar unidentifiability issue may arise in SSEs. Responding to this concern, a recent study mathematically confirmed the identifiability of a specific subset of pure-birth SSE models (16). Meanwhile, another study stated that SSEs are identifiable (17).

Unlike previous studies, we demonstrate that model congruence is a universal phenomenon across all SSEs using the general model HiClaSSE, which incorporates common SSEs as special cases (e.g., BiSSE, MuSSE, HiSSE, MiSSE, BiSSE-ness, and ClaSSE). This congruence arises from the hidden states in SSEs and the lumpability properties of Markov models (18). As a result, an SSE, whether it includes hidden states or not, is congruent with an infinite set of models that do include hidden states. These models form a congruence class in which SSEs have different diversification histories and parameter counts. For example, any state-independent model is congruent with trait-dependent models that have the same or fewer parameters.

Consequently, this congruence poses a challenge for testing trait-dependent diversification. We demonstrate that traditional model selection methods fail to identify the correct trait-diversification scenario within the same congruence class. To address this, we propose an analytical solution that selects the simplest model within a congruence class, without significantly altering existing phylogenetic practices. Additionally, we show that the discovered congruence is the only type of congruence possible in SSEs. Thus, our study highlights that model unidentifiability is not harmful to macroevolutionary inference with SSEs.

However, we find that traditional model selection approaches consistently fail to select the correct traitdiversification scenario across different congruence classes, with an extremely high false positive rate. SSEs have long been known to be susceptible to this issue (3). Interestingly, the discovered congruence provides a clear explanation for this problem. While we do not have a definitive solution, we discuss potential approaches below. Our conclusions are supported by both simulations and empirical data.

## Results

### HiClaSSE and Hidden States

The HiClaSSE model extends ClaSSE (19) and BiSSE-ness (20) by including hidden states, as in HiSSE (5). In addition to speciation and extinction events (1), HiClaSSE allows modeling two types of state-change events. The cladogenetic events which denote state transitions occurring with lineage bifurcation during speciation, and anagenetic events which denote state transitions without speciation. For simplicity, speciation events with and without state changes are termed coupled and decoupled speciations, respectively. HiClaSSE may include any number of states *k*.

Like other SSEs, HiClaSSE is a joint process that combines an observable trait and a diversification component. The trait represents a Continuous Time Markov Chain (CTMC) where discrete states change over time. The diversification component is a separate CTMC whose states are unobservable diversification regimes, each equipped with a specific speciation and extinction rate to control lineage birth and death. Thus, in an SSE, every state is effectively hidden, belonging simultaneously to the trait state and the diversification regime. Observable states are modeled at the tips of tree using ambiguous character coding, they are the collections of the hidden states.

For example, in a four-state HiSSE that characterizes dependent evolution between a binary trait and a diversification process with two regimes each observable state comprises two hidden states. Conversely, a trait-free MiSSE (6), which models diversification shifts alone, maps all hidden states onto a single observable one; thus, the trait CTMC disappears in MiSSE, leaving only the CTMC for diversification regimes. This framework allows for the construction of various SSE models. Note, our interpretation of hidden states, for biological clarity and mathematical precision, may differ from previous treatments (SI Appendix).

### HiClaSSE Likelihood

HiClaSSE likelihood is calculated by solving a system of ODEs backward in time using two terms: ***D***_***i***_(*t*), the probability of a lineage being in state *i* at time *t*, and *E*_*i*_(*t*), the probability of a lineage in state *i* not surviving until the present time (1, 21). Representing them as column vectors, ***D*** = [*D*_1_(*t*),…, *D*_*k*_(*t*)] and ***E*** = [*E*_1_(*t*),…, *E*_*k*_(*t*)], yields the following ODEs expressed in array notation:

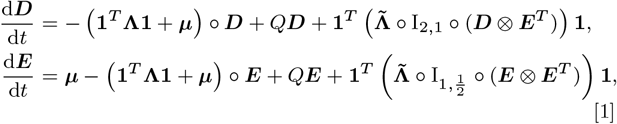

where, “○” is an element-wise product of two arrays, ***D****⨂****E***^*T*^ is outer product of two vectors, and I_*a,b*_ indicates square matrix where all diagonal elements are *a*’ s while off-diagonals are *b*’s. The speciation tensor **A** = {Λ_1_,…, Λ_*k*_} contains *k* × *k* matrices Λ_*i*_ with speciation rates (λ_*ijz*_). For example, the notation *λ*_111_ indicates a decoupled speciation event – when one lineage in state 1 speciates into two lineages with the same state. In contrast, the notationλ_122_ indicates a symmetric coupled speciation – when one lineage in state 1 produces two lineages each with the state 2. Similarly, λ_123_ denotes an asymmetric coupled speciation (19, 21).

The extinction rates of a lineage are denoted by the vector ***µ*** = [*µ*_1_(*t*),…, *µ*_*k*_(*t*)]. The instantaneous rate matrix *Q* defines anagenetic transitions (*q*_*ij*_) within a CTMC that governs the joint evolution of the trait states and diversification regimes (details in SI Appendix). HiClaSSE is a general model that includes various SSEs as submodels. For instance, HiClaSSE with two states and no cladogenetic events is a BiSSE model (1). HiClaSSE is similar to SecSSE (22), another general SSE model that differs in how it conditions the overall likelihood at the root on clade survival. All findings in this paper apply to SecSSE as well.

### Correlated Evolution in SSEs

In SSEs, the parameters determine the relationships between the trait and the diversification process. Trait-dependence occurs when: (**C1**) there is variation in speciation and/or extinction rates between observable states as in BiSSE (1); or (**C2**) a model includes cladogenetic events that result in shifts of diversification regimes as in ClaSSE and BiSSE-ness (19, 20). However, this list is not exhaustive. It is often overlooked that SSEs with hidden states may have additional parameterizations implying trait-dependence. While these parameterizations were initially formulated for binary traits (23), they can be readily extended to SSEs. Specifically, they are applicable when the condition (**C1**) is met, and they arise in the following situations: (**C3**) when the rate of change in the diversification regime depends on the trait; (**C4**) when changes in both the trait and regime are interdependent; and (**C5**) when changes in the trait and regime occur simultaneously, typically when dual state transitions are allowed (SI Appendix).

It is important to note that conditions (**C3-5**) imply that trait-dependent diversification does not necessarily require a distinct set of speciation or extinction rates for each observable state. It can also arise from differential transition rates between hidden states in a SSE (e.g., Fig. 5C-F, E).

### Lumpability and Congruence

Modeling heterogeneity in the diversification process often requires hidden states. We find that hidden states can lead to congruent models. This congruence arises from the principle of strong lumpability (18, 24), which we refer to simply as “lumpability”. Lumpability occurs when the hidden states in an original SSE (*S*_1_) can be reduced through aggregation, resulting in a model with fewer states (*S*_2_), making both models congruent (*S*_1_ ≅*S*_2_, SI Appendix). Note that this aggregation must preserve the number of observable states. Lumpability depends on specific rate symmetries in the original SSE. Without these symmetries, aggregation and congruence are not possible, and a simpler SSE with fewer states cannot exist.

**Table 1.** Results of SSE models for the Phasmatodea dataset.

| Model | np | $\cong$ | Trait-dep. | Ln | AIC | ΔAIC |
| --- | --- | --- | --- | --- | --- | --- |
| CID4 | 5 | – | no | -988.22 | 1986.44 | 10.02 |
| CID2 (BiSSE) | 4 | no | no | -991.05 | 1990.09 | 13.68 |
| CID8 | 5 | yes | no | -988.22 | 1986.44 | 10.02 |
| COR8-C | 5 | yes | yes | -988.22 | 1986.44 | 10.02 |
| CLA8-C | 5 | yes | yes | -988.22 | 1986.44 | 10.02 |
| EHE25-C | 3 | yes* | yes | -988.28 | 1982.57 | 6.15 |
| COR8-NC | 5 | no | yes | -988.21 | 1986.42 | 10.00 |
| CLA8-NC | 5 | no | yes | -992.18 | 1994.36 | 17.94 |
| **SEM8** | **4** | **no** | **yes** | **-984.21** | **1976.42** | **0.00** |

To identify these rate symmetries, we derived the exact lumpability conditions for SSEs by applying the approach for standard CTMCs (18) to the SSE’s likelihood in Eq. 1 (SI Appendix).

Specifically, an SSE model is lumpable if three following conditions are met simultaneously (SI Appendix): (**L1**) the rate matrix *Q* is lumpable, which implies that for any two aggregated states *ŝ*_*i*_ and *ŝ*_*j*_, the total transition rate 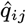, from any original state *s*_*m*_ in *ŝ*_*i*_ to *ŝ*_*j*_ remains constant. These 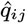 ‘s become the transition rates in the lumped *Q* (18); (**L2**) all *µ*’s within each partition set must be equal, and the lumped extinction rate is that value; (**L3**) the speciation tensor presents a more complex situation. In this case, each lumped speciation rate 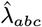 is composed of the original rates *λ*_*ijz*_ which can be grouped based on the parent branch state (i.e., *i*). So, the speciation tensor is lumpable if the sum of the original λ’s in each group is the same. The lumped speciation rate is one sum that satisfies this condition.

To investigate congruence, we can construct congruent models in reverse using an operation called “hidden expansion” (HE). This involves adding hidden states to an initial model *S*_0_ to create a new model *S*_1_ that maintains lumpability with *S*_0_, implying that *S*_0_ ≅ *S*_1_.

As an example, consider the trait-free MiSSE model (*M*_1_ in Eq. 2) with three parameters (*λ*_*AAA*_, *λ*_*BBB*_, *q, µ*). This model is congruent with another model, *M*_2_ (Eq. 2), in which there are three diversification regimes, but the speciation rates for the regimes *B* and *C* are the same (i.e., *λ*_*BBB*_ = *λ*_*CCC*_). Clearly, *M*_2_ is lumpable given the state partition {{*A*}, {*B, C*}} as it meets the lumpability conditions (**L1-3**). Since all states in *M*_2_ are hidden, it collapses to *M*_1_ under the lumping.

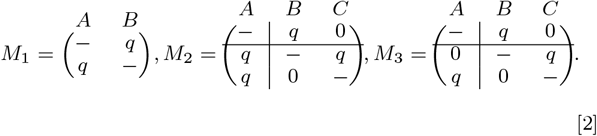

In contrast, a slightly different model *M*_3_ (Eq. 2) is not congruent with either *M*_1_ or *M*_2_, as its rate matrix violates the lumpability condition (**L1**) for the partition {{*A*}, {*B, C*}} because the transitions *B* ⟶ *A* and *C ⟶ A* are not equal. Therefore, despite all three models have the same parameter count, *M*_3_ stands as a distinct model. Note, the states of *M*_3_ cannot be reduced to a two-state model. While sharing the same speciation rates as *M*_1_, *M*_3_ exhibits distinct diversification dynamics, which leads to a different pattern of speciation and extinction events on a phylogeny. This result is supported by our simulations yielding identical likelihoods for *M*_1_ and *M*_2_.

### Hidden Expansion and Congruence Class

While the example above illustrates a fairly trivial HE, this technique can be universally applied to any SSE to generate congruent models. For instance, *M*_2_ can be further expanded, maintaining lumpability, to create models with any number of hidden states. With limitless possibilities for preserving lumpability, countless implementations of HE exist. Thus, every SSE model belongs to an infinite congruence class, a phenomenon we define as HE-congruence.

### Congruence between CID and Dependent SSEs

Any traitindependent model (CID) is congruent with an infinite set of trait-dependent SSEs. For example, CID4, with four parameters (Fig. 5A: *λ*_1_, *λ*_2_, *q, µ*), can be expanded into a congruent CID8 model (Fig. 5B) by adding four hidden states (SI Appendix). Note, the numeral in the model denotes the count of hidden states. Rewiring the rates (*q*’s) and (*λ*’s) in CID8, while keeping the model lumpable with CID4, yields congruent trait-dependent models like COR8-C and CLA8-C (Fig. 5C-D). This congruence arises purely from lumpability and is supported by our simulations, which yielded identical likelihoods for CID4 and its congruent counterparts. This method typically preserves the parameter count; the resulting congruence class is infinite – for example, CID4 can be expanded to CID16 with similar changes.

### Equal Rate Hidden Expansion (EHE)

HE can be extended to a point when all permissible rates are equal, except extinctions. We refer to this transformation as Equal Rate Hidden Expansion (EHE). Consider, for instance, a CID4 model with four parameters: *λ*_1_ = 0.3, *λ*_2_ = 0.1, *q* = 0.2, and constant *µ* (Fig. 5H). Applying the EHE to this model results in the EHE8-C (Fig. 5I), where the diversification is trait-dependent (due to **C2, C3, C5**), and includes coupled speciation events. The EHE8-C has only two parameters compared to the four in the CID4, but all its *λ*’s and *q*’s have the same value of 0.1, while extinction remains unchanged (SI Appendix). Our simulations confirm that CID4 and EHE8-C yield identical likelihoods.

The EHE transformation reduces the number of parameters to the number of extinction categories plus one and can be applied to any SSE, regardless of state count or rate values. This universality stems from the properties of rational numbers, because any rational can be expressed as the product of an integer *z* and a rational multiplier *r*. If the rates are not rational, they can be approximated to the desired precision. As a result, the set of parameters (*θ*’s) in an original SSE, excluding extinction, expands as {*θ*_1_ = *z*_1_*r, θ*_2_ = *z*_2_*r*, … *θ*_*n*_ = *z*_*n*_*r*}. The factor *r* serves as the common rate, while the integers *z*_*i*_ roughly correspond to the number of additional hidden states. An SSE that is sufficiently large to embed these additional states represents the EHE transformation. We present a general algorithm that can be used to create EHE models (SI Appendix).

For a given SSE model, EHE generates an infinite congruence class of trait-dependent models because there is an infinite set of multipliers *r* to use for the same original model. The choice of *r* determines the state count in the EHE model. Typically, EHEs with more states provide a better approximation of the original model. A universal parametric form of EHE does not exist because it depends on the rate values in the original SSE (SI Appendix). For example, varying parameter values in the CID4 (Fig. 5H) would result in different EHE models.

### Semi-Congruent Models

HE can also generate trait-dependent models from different congruence classes with a reduced parameter count. For example, adding four hidden states to CID4 (Fig. 5A) and rewiring the rates without adhering to the lumpability conditions may result in a SEM8 model (Fig. 5E). We refer to such irreducible models as semi-congruent (SEM) when they have fewer parameters but still resemble the original model. For instance, SEM8 (Fig. 5E) has fewer parameters than CID4 (Fig. 5A) but remains topologically similar to EHE models within the CID4 congruence class (e.g., EHE8-C in Fig. 5I). Unlike EHE models, SEM require only a few extra hidden states.

SEM models can be generated in various ways and may provide a better fit than the true model. To explore this, we developed a simple algorithm that traverses the space of 531,441 possible models by successively testing only 36 different transition rate combinations in *Q* to save computational time. While not the most efficient, it serves our purpose. We simulated data under CID4 with four parameters (Fig. 5H) and applied our algorithm to each dataset to search for the best SEM8 constrained to just two parameters. In 81-91% of simulations, trait-dependent SEM8 was preferred over CID4 (6-AIC > 2), providing a substantially better AIC fit overall (Fig. 1). When using the Bayesian Information Criterion (BIC), SEM8 was selected in every trial (6-BIC > 2, Fig. 1). Despite having fewer parameters, SEM8 also yielded higher likelihood scores in 56-71% of trials, depending on the number of tips.

**Fig 1.**
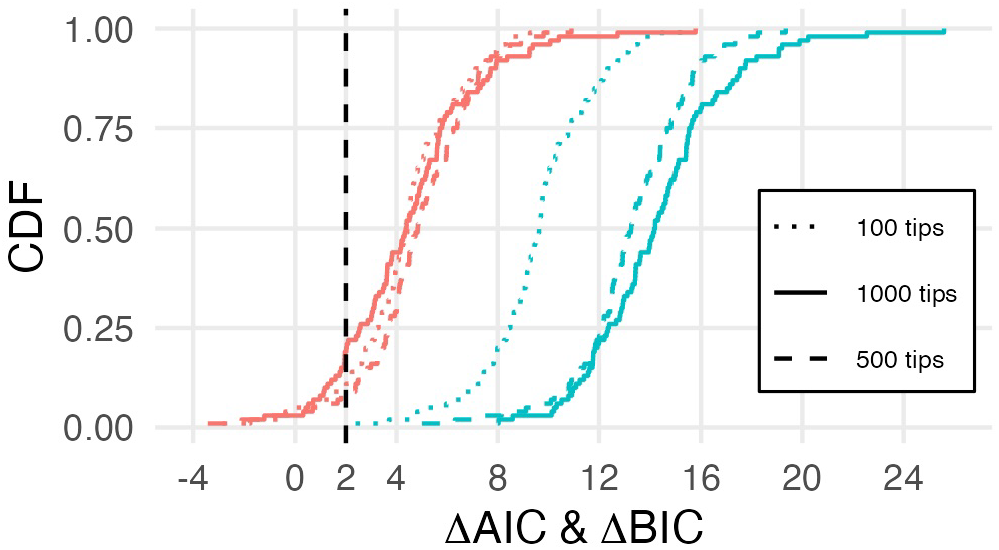
Performance of semi-congruent models illustrated using the cumulative distribution function (CDF) of .6.AIC (red) and .6.BIC (blue). Positive values indicate support for trait-dependent semi-congruent models (SEM8) with two parameters, compared to the true CID4 with four parameters. SEM8 and CID4 belong to different congruence classes. Simulations were conducted on phylogenetic trees with 100, 500, and 1000 tips.

### The Structure of a Congruence Class

Each congruence class, generated by HE, consists of a unique non-lumpable model called the irreducible model (*M*_0_) and an infinite set of lumpable models created by adding hidden states to *M*_0_ (Fig. 2). Thus, all other models in the class have more hidden states than *M*_0_. For instance, CID4 and BiSSE are irreducible models with and without hidden states, respectively. In addition to the HE explained earlier, which lead to congruent models with the same or fewer parameters, it is straightforward to create models with more parameters.

**Fig 2.**
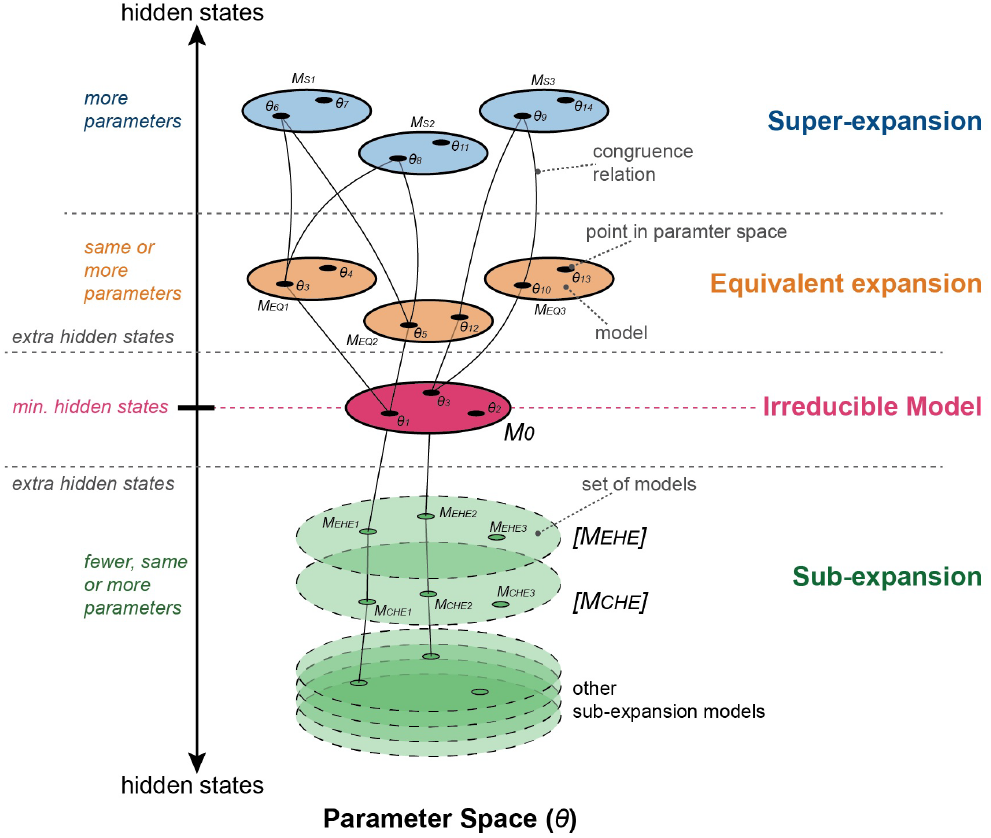
Structure of HE-congruence class. The irreducible model *M*_0_ is congruent with all equivalent and super-expansion models within the class (e.g, *M*_0_ *≅ M*_*EQ*1_ *≅ M*_*S*1_). Only specific parameter values of *M*_0_ are congruent with sub-expansion models (e.g, *M*_0_ (*θ* _1_) ≅ *M*_*EHE*1_), while the entire *M*_0_ is congruent with a set of sub-expansion models (e.g, *M*_0_ *≅* [*M*_*EHE*_ ]). Note, congruence relationships are transitive, not all of them are shown for clarity; * denotes the parameter count for off-diagonal parameters in *Q* (SI Appendix).

We identified three main types of HE: sub-expansion (e.g,. EHE), equivalent (e.g, CID8, COR8-C, CLA8-C), and super-expansion. These types create models with the fewer, same, or more parameters in the off-diagonal blocks of the *Q* matrix, respectively. The off-diagonal parameters are essential for defining the parametric form (SI Appendix).

Sub-expansion models depend on specific parameter values in *M*_0_. Each parameter setup corresponds to a distinct sub-expansion model, usually with fewer parameters than *M*_0_. *M*_0_ is not directly congruent with individual sub-expansion models but rather with sets of these models (e.g., *M*_0_ ≅ [*EHE*]).

### Topological bridges

The probabilistic behavior of SSEs does not necessarily align with their classification as trait-dependent or independent. This mismatch arises due to topological bridges created by congruence classes in the SSE model space. For example, CID4 and COR8-NC (Fig. 5A,F) differ in their parameters and state counts, yet their probabilistic behaviors are similar because CID4’s congruence class includes COR8-C (Fig. 5C), which is topologically close to COR8-NC. As a result, data generated by CID4 may fit the trait-dependent COR8-NC model better than other irreducible CID models from different congruence classes (Fig. 3).

**Fig 3.**
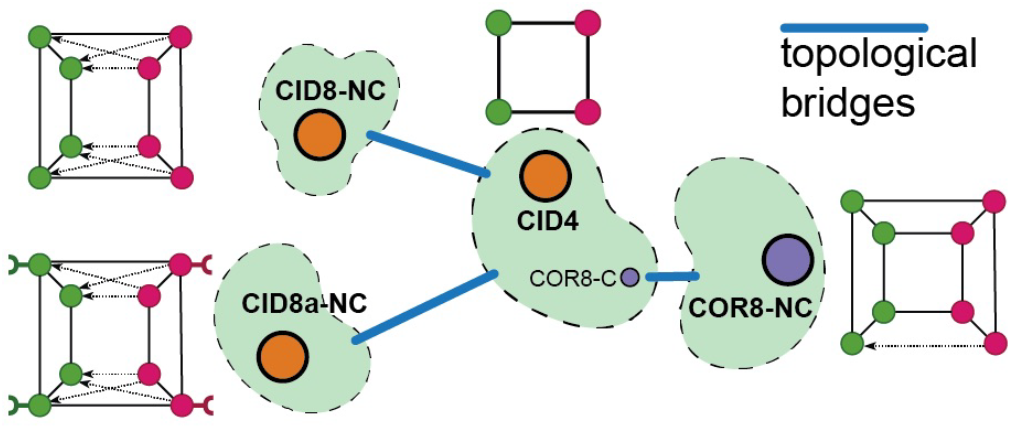
Congruence classes (in green) create topological bridges in the SSE model space, making CID4 probabilistically closer to some irreducible trait-dependent SSEs (e.g., COR8-NC) than to other irreducible CID models, such as CID8-NC with unidirectional transitions and CID8a-NC with coupled speciation rates. Note that all models have the same parameter count; see the text and Fig. 5 for details.

Our simulations with expected Δ*AIC* support this finding. The expected Δ*AIC* is a proxy for relative Kullback-Leibler (KL) divergence, which measures the similarities between two probability distributions (25). The expected Δ*AIC* over 100 trials for data generated under CID4 was 0.4, 2, and 5 for COR8-NC, CID8-NC, and CID8a-NC, respectively. Moreover, this phenomenon becomes more pronounced with increasing amounts of observed data. In this setup, excluding the true CID4 from the analysis may lead to a false positive selection of the trait-dependent scenario. This phenomenon applies to all SSEs, as each has congruent models with hidden states.

### Other Congruence Requires Weak Lumpability

Consider two HiClaSSE models, *H*_0_ and *G*_0_, from different HE-congruence classes. Can another type of congruence exist between them? Assume that both *H*_0_ and *G*_0_ are irreducible, meaning each has an HE-congruence class that includes SSEs with more hidden states (Fig. 4). There always exists a model *G*_*s*_ such that *G*_*s*_ *≅ G*_0_, and *G*_*s*_ has more hidden states than *H*_0_. If *H*_0_ is congruent to *G*_0_ through another type of congruence, then *H*_0_ must also be congruent to *G*_*s*_. This suggests that *G*_*s*_ (with more hidden states) can be aggregated into *H*_0_ (with fewer states), but this aggregation does not represent strong lumpability since *G*_*s*_ and *H*_0_ belong to different HE-congruence classes. Interestingly, this type of aggregation is known as weak lumpability (18, 26). It occurs when models are still lumpable, but the conditions for strong lumpability (**L1-L3**) are not satisfied. Unlike strong lumpability, which holds under any initial probability vector at the tree root, weak lumpability depends on a specific vector. Therefore, the existence of other types of congruence in HiClaSSE necessarily requires weak lumpability.

**Fig 4.**
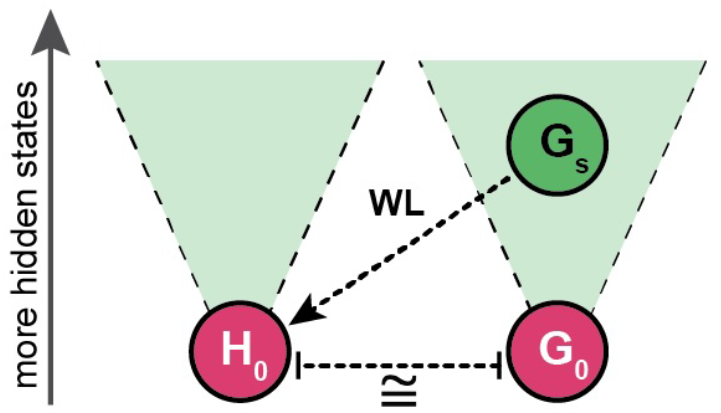
Tow irreducible models *H*_0_ and *G*_0_ and their HE-congruence classes (in green). Assuming that *H*_0_ and *G*_0_ are congruent through some other type of congruence (non-HE), would imply that *G*_*s*_ is weakly lumpable (WL) into *H*_0_, where *G*_*s*_ *≅ G*_0_ and *G*_*s*_ has more hidden states than *H*_0_.

**Fig 5.**
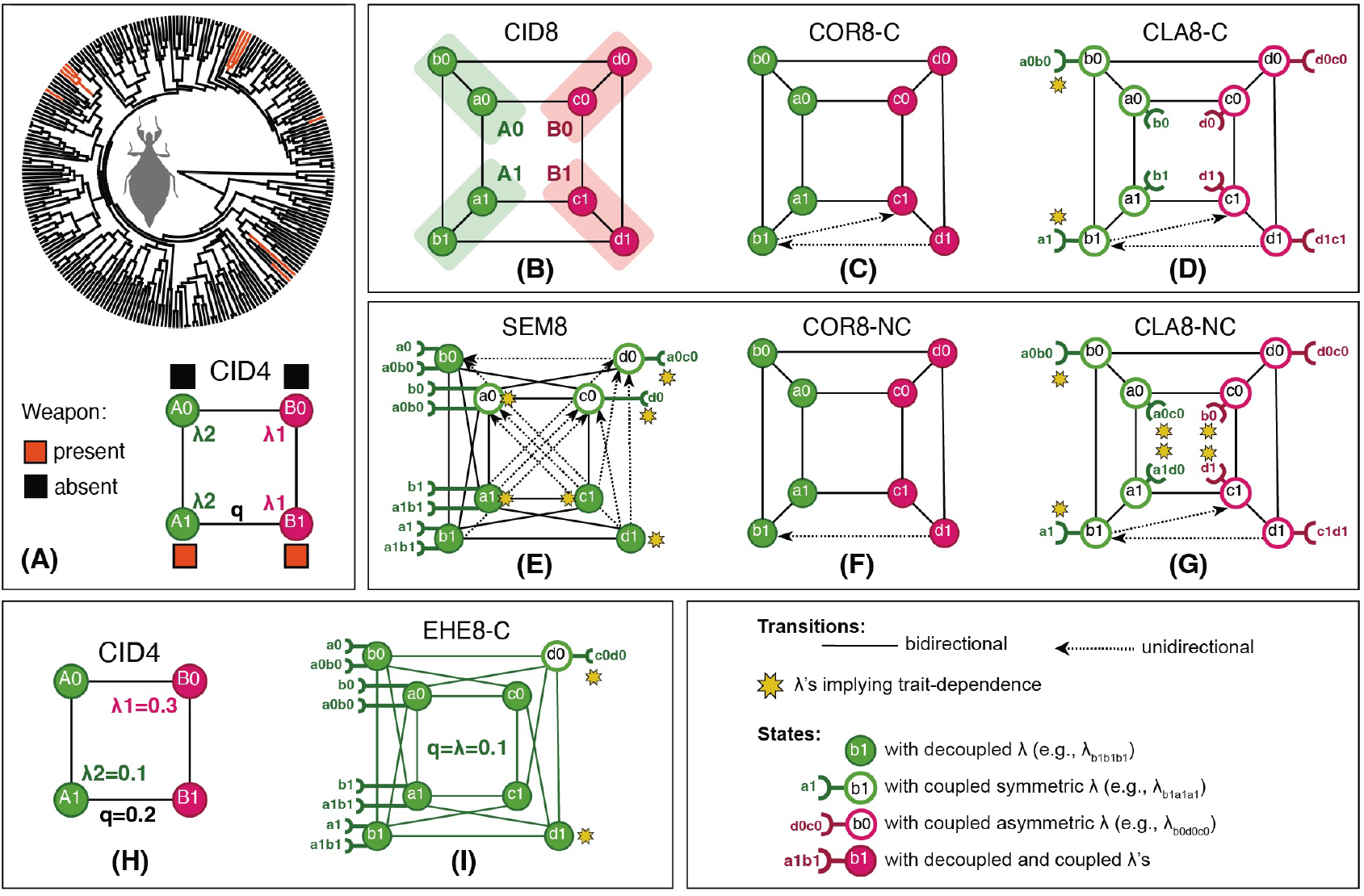
SSEs for the Phasmatodea dataset with 243 tips (A-G, Table 1), and simulations (B-D, F-G, H-I); see text for details. For Phasmatodea, the SSEs include two extinction rates, while for the simulations one extinction rate. Different colors in the model diagrams represent distinct rate categories. All models are derived from CID4 through hidden expansion; (B) shows the mapping of states between CID4 and the expanded models. (A-B, H) Trait-independent SSEs. (C-G, I) Trait-dependent SSEs. Cause of dependence: (C-F) due to condition (**C3**); (D-G) due to conditions (**C2**,**3**); and (E-I) due to conditions (**C2**,**3**,**5**). (B-D) Congruent to CID4 in (A). (I) Congruent to CID4 in (H). (E-G) Non-congruent to CID4.

## Discussion

Using the general HiClaSSE model, we demonstrated that every SSE belongs to a congruence class that includes an infinite set of models producing identical likelihoods, regardless of the amount of data. Thus, congruence is ubiquitous in SSEs, which contradicts the previous study claiming that SSEs are identifiable (17), as it did not consider hidden expansion (HE).

Congruence arises through HE by adding hidden states to a model, while preserving the lumpability conditions in the expanded model, resulting in both models being congruent. Thus, lumpability helps differentiate between SSEs that appear distinct but are fundamentally identical (i.e., congruent), and those that are genuinely different (i.e., non-congruent). Congruence holds under any sampling fraction of extant species (SI Appendix). HE congruence classes are akin to those identified by LP for time-dependent diversification models (7), due to rate invariance. In HE classes, the invariance is reflected in the distribution of rates across observable states, while in LP classes, it is associated with pulled speciation and diversification rates.

Models within the same congruence class share identical extinction parameters but differ in the number of speciation and transition parameters, ranging from one (sub-expansions) to infinity (super-expansions). Specifically, any trait-independent model (e.g., CID) is congruent with an infinite set of trait-dependent models that have the same or fewer parameters. Since congruence is symmetric, this also implies that some trait-dependent models are indistinguishable from trait-independent models due to unidentifiability.

This raises four important questions: does congruence mislead hypothesis testing in trait-diversification relationships? Can we reliably select the correct model both within and across congruence classes? How might this impact current phylogenetic practices? And what insights can we gain to enhance existing practices?

### Selecting within Congruence Classes

The standard practice for testing trait-dependent diversification involves fitting alternative SSEs corresponding to trait-independent and -dependent scenarios and then assessing their fit using, for example, ΔAIC (4).

A recent study on stick insects (Phasmatodea), where males use leg protrusions as weapons in combat for females, found no effect of this trait on diversification (27). The study tested five alternative models, ranging from BiSSE to HiSSE and CID. The best-fitting model was CID4 (Fig. 5A), which outperformed the closest trait-dependent model (HiSSE) by ΔAIC = 27.7.

The theory outlined above suggests that there are myriads of scenarios congruent with the identified CID4. While some are equally trait-independent (e.g., CID8, Fig. 5B), others imply trait-dependent diversification. They range from scenarios where the weapon affects speciation by altering transitions between diversification regimes (COR8-C, Fig. 5C), to more complex dependencies. For example, in CLA8-C (Fig. 5D), diversification regimes shift at speciation events, which are dependent on the weapon trait (indicated by stars in Fig. 5D), with all speciation events coupled with state changes. These models have the same parameter count as CID4, and hence yield identical AIC scores (Table 1). Since CID4 is the null model and the AIC scores are the same, CID4 remains preferred, meaning these alternatives do not affect model selection. Note that congruence arises from lumpability; slightly breaking lumpability yields trait-dependent models from different congruence classes (e.g., COR8-NC, CLA8-NC, Fig. 5F-G, Table 1).

Yet, other congruent but trait-dependent scenarios can be created using Equal Rate Hidden Expansion (EHE) models, which have fewer parameters – specifically, the number of extinction rate categories plus one. In EHEs, certain diversification shifts, coupled with speciation events, are trait-dependent (e.g., indicated by star in Fig. 5I). These models typically involve thousands of hidden states, making them computationally impractical. In this example, we use an EHE model with 25 hidden states (EHE25-C), which approximates CID4’s rates to three decimal places. While increasing the hidden states can achieve any desired approximation, this is unnecessary as our analytical approach derives them without fitting directly to data (SI Appendix). Notably, AIC selects EHE25-C over CID4 due to fewer parameters (Table 1). Since there is an infinite number of EHEs in any congruent class, this implies that an infinite number of trait-dependent models exist for any CID.

The overviewed congruent SSEs are not part of the standard set, yet they represent plausible scenarios with varying numbers of trait-dependent anagenetic and cladogenetic events, which cannot be ignored. Does this mean we should accept the trait-dependent scenario suggested by EHE25-C? Both theory and simulations show that the same AIC performance manifests in any congruent class of a CID model. The Phasmatodea example itself highlights why the trait-dependent scenario should be questioned – the data are simply limited, consisting of only 10 tips with the weapon trait evolving six independent times. Reliable inference with SSEs heavily depends on sufficient character sampling (28). Thus, we agree with the original study and consider CID4 should remain the preferred model. Given this evidence, it is obvious that AIC misleads model selection within a congruence class, as it does not penalize model complexity for increasing numbers of hidden states.

### Solution for Within-Class Selection

Our results indicate that a simple parameter count in SSEs fails to adequately capture model complexity, as has been observed in non-phylogenetic hidden Markov models (29). Thus, hidden states tend to substitute for individual parameters in SSEs, making AIC generally misleading, especially when the hidden state count varies across candidate models. EHEs, where parameter complexity is largely replaced by complexity in the topology of hidden states, illustrate this phenomenon well, highlighting why AIC results should not be blindly followed. We do not anticipate any statistically valid correction to AIC that adds a penalty as a function of hidden state count. Moreover, none of the current software (e.g., the HiSSE R package) includes such corrections.

We propose a solution by applying the parsimony principle when selecting among congruent models, favoring a model that explains rate variation with the minimal number of hidden states. This approach leads to a unique solution for each congruence class, known as the irreducible model *M*_0_ (Fig. 2), which can be derived analytically by lumping the states in a reducible model. In the stick insect example, the CID4 is the irreducible model, with all other congruent models collapsing into it. Generally, this method consistently favors CID models over others within their congruent classes.

Our parsimony principle is primarily a practical assumption for hypothesis testing, introducing the minimum set of restrictions, as the uniqueness of *M*_0_ renders model selection unambiguous. However, we acknowledge that in some cases, other models within the same class may more accurately reflect the true diversification process, but they remain unidentifiable.

Our main recommendation is to always use irreducible models for inference to avoid unidentifiability issues. This is straightforward if a model has no hidden states, as all such models are irreducible. We did not identify any reducible models among the standard SSEs provided by RevBayes (21, 30) or HiSSE (5). However, congruent models can still be constructed using these software. The proposed solution suggests that, in general, previous studies do not require reevaluation in light of identified congruence, but users should remain vigilant about this pitfall.

### Selecting across Congruence Classes

SSEs are prone to false positives, or Type 1 errors, where a true null hypothesis is incorrectly rejected. These errors often result from model misspecification, where the null model fails to adequately capture the complexity of the underlying diversification process (3). Apparently, minimizing Type 1 errors can be achieved by testing a broader set of alternative models (4, 5) from different congruence classes. However, this approach will only work if AIC can accurately select SSEs across congruence classes, without being subject to the same bias present within a congruence class.

We simulated this scenario by testing the performance of the true CID4 against a pool of ≥0.5M candidate SSEs using the optimization algorithm designed to find the best-fitting model. Surprisingly, AIC consistently selected incorrect trait-dependent models with fewer parameters (SEM8) over CID4, with strong support (Fig. 1). Even for models with eight states (e.g., SEM8), the space of possible parameterizations is too vast (31) to be efficiently traversed using our algorithm. We expect that exploring a broader range of models will further increase the Type 1 error rate. This result suggests that it is almost always possible to find a false SSE that fits the data better than the true one. Thus, AIC fails to select the appropriate model if the candidate pool is large enough.

Applying the same approach to Phasmatodea yielded a similar result, with strong support for the irreducible trait-dependent model (SEM8, Fig. 5E), which significantly out-performed CID4 and EHE25-C, with ΔAIC = 10.02 and 6.15, respectively (Table 1). In SEM8, the weapon trait causes diversification regimes to evolve anagenetically rather than cladogenetically, and affects transitions between the regimes. Given the sampling limitations in the Phasmatodea dataset, and the fact that simulations corroborate the empirical data, we consider this result to be a false positive too, rather than evidence of trait-dependent diversification.

### Model Proliferation and Misspecification

We found that the primary cause of AIC’s misestimation of model complexity stems from the vast diversity of possible SSEs that can be constructed to model trait-diversification processes. This diversity leads to model “proliferation”, where an incorrect model from a large pool fits the data better than the true model simply by chance. In essence, model proliferation is akin to the multiple comparisons problem in statistics (32), where the false positive rate increases with the number of comparisons, even if the rate is low for each comparison. Consequently, as the pool of candidate SSEs grows, the risk of Type 1 error rises accordingly.

Our semi-congruent simulations (Fig. 1) support the model proliferation problem: in almost every simulated dataset, a different false model (SEM8) was preferred, with the proportion of unique SEM8s ranging from 81-91% across trials, depending on the number of tips.

Importantly, in both within- and across-congruence class cases, the misestimation of complexity stems from model proliferation. Within a class, this misestimation is clearly revealed by analytically available congruent models (e.g., EHEs), while across classes, it requires computational exploration. For a given number of states, there are more trait-dependent than -independent SSEs (31). Thus, for each dataset generated under CID, there is almost always an irreducible and better-fitting trait-dependent model due to proliferation (Fig. 6A). This source of Type 1 error contrasts with model misspecification, which occurs when the adequate model is not included in the analysis (Fig. 6B).

**Fig 6.**
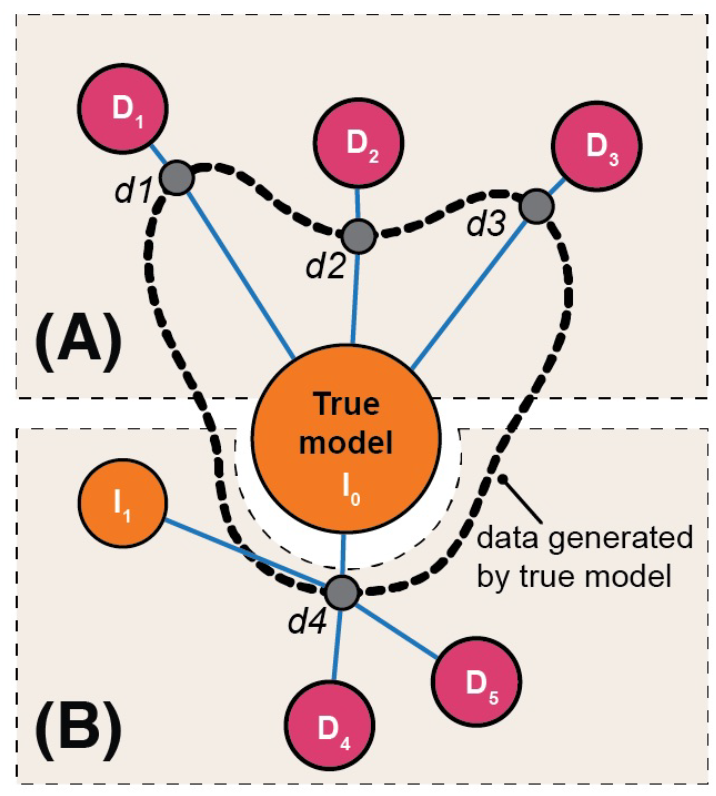
Scenarios of model (A) proliferation and (B) misspecification, leading to Type 1 errors. Trait-dependent and trait-independent models are represented by red and orange circles, respectively; a better model fit is indicated by shorter blue paths. In (A), the true CID (*I*_0_) is included in the analysis, but for each possible dataset (*d*_1_ -*d*_3_), a separate better-fitting SSE (*D*_1_ -*D*_3_) can be found purely by chance, each fitting only its specific dataset. In (B), the true model (*I*_0_) is missing from the analysis; although another CID (*I*_1_) is included, it fits worse than the alternative models (*D*_4_ -*D*_5_) due to model inadequacy.

Our theoretical findings help explain why misspecification is such a common issue in SSEs. Congruence classes create topological bridges in model space, where the probabilistic behavior of a CID may be closer to trait-dependent models than to other CIDs (Fig. 3). As a result, the set of all CIDs does not form a distinct model “family” within SSEs; it would if, for instance, all CIDs were nested within each other and within dependent SSEs. This configuration violates the fundamental assumption of the Maximum Likelihood (ML) estimator, which requires the true data-generating process to be within the the family of models used for inference (33). Thus, the magnitude of model misspecification should not be surprising. When an adequate CID model is missing from the candidate set, a dependent model – though incorrect – may appear to fit the data better than the chosen CIDs, due to topological bridges (Fig. 6B). A similar issue has been observed in diversity-dependent birth-death models (34).

Our findings suggest that the problem is even more severe. To reduce the risk of misspecification and account for the complexity of real evolutionary process, a larger pool of candidates must be tested. However, this rapidly increases the risk of model proliferation: the vast diversity of possible SSEs makes it likely that the wrong model will be selected purely by chance, even if the true model is present.

### Implications

We have demonstrated that AIC consistently fails in hypothesis testing both within and across congruence classes. However, within a single congruence class this issue is not problematic when applying the proposed parsimony refinement. The challenge arises when selecting models across different congruence classes due to the interference of model proliferation and misspecification, leading to an extremely high rate of Type 1 errors. Applying other traditional criteria, such as Akaike weights or BIC, will not help, as they count parameters in a similar way. In fact, BIC resulted in an even higher false positive rate, reaching 100% in simulations (Fig. 1). In contrast, Bayes Factor (BF) requires separate investigation, as it integrates over prior space, including priors on tip states, which are highly uncertain in hidden state models. This uncertainty may be treated as a “parameter” by BF, potentially penalizing complex models.

A robust solution for model selection across congruence classes is urgently needed, as it will make SSEs with hidden states a powerful tool for detecting trait-diversification relationships. In the meantime, we recommend exercising caution when using SSEs for hypothesis testing. To minimize Type 1 errors, we suggest the following: (1) test various null CID models from different congruence classes; (2) consider constraining models using prior biological knowledge (8); (3) rely only on substantial AIC scores as evidence for trait-dependent scenario; (4) use sufficiently large phylogenies with representative sampling of character states at the tips (28); and (5) ensure that trait-dependent scenarios are supported by reasonable biological interpretations. Additionally, users may adapt the statistical solutions discussed in the following section to improve their analyses. It is important to note that congruence also holds in models for ancestral state reconstruction alone (i.e., CTMCs), as they are submodels of SSEs without diversification.

Our study emphasizes that the concept of lumpability is an essential tool for investigating SSE behavior, just as it is for CTMCs alone (24, 35). As we have demonstrated, lumpability is critical to understanding the relationships between SSEs and precisely explaining the sources of Type 1 error. Additionally, lumpability uncovers a wide range of unexplored CID models, which can be created by combining various irreducible MiSSEs with irreducible trait CTMCs into a joint process (e.g., Fig. 3). These SSEs may be promising for better modeling the complexity of biologically-realistic trait-diversification process (36).

### Ways forward

To minimize model misspecification, the possible solution must efficiently explore the model space. In a Bayesian framework (BI), reversible-jump MCMC has been shown to be effective for CTMCs, which are submodels of SSEs (31). However, the key challenge lies in designing efficient trans-dimensional moves to traverse across hidden expansions (37). In the ML framework, optimization algorithms may serve as a solution by navigating through hidden expansions and optimizing model fit.

The optimization algorithms will be highly susceptible to model proliferation, and this issue may persist in reversible-jump MCMC. In BI, applying priors on the number of hidden states can mitigate this effect. Since model proliferation occurs largely due to chance, false-selected models should exhibit low predictive power. Consequently, Bayesian posterior prediction and parametric bootstrapping in ML offer promising alternatives (38, 39). Additionally, cross-validation techniques, applied in both BI and ML to assess a model’s predictive ability, show potential (39, 40). These techniques typically split the data into training and validation sets; however, it’s unclear how they would perform on datasets influenced by singular rate-shift events, which may be unevenly distributed across the splits. The primary challenge for both posterior prediction and cross-validation lies in combining these methods with efficient exploration of the model space.

### Other Congruence Classes?

We showed that in time-independent SSEs (=HiClaSSE), the existence of any type of congruence other than HE-congruence would inevitably require weak lumpability (Fig. 4), even if it is not associated with hidden states. Since weak lumpability does not hold on phylogenetic trees (24), this rules out the existence of other congruence classes. This is also supported by the fact that pure-birth SSEs are identifiable (16). Consequently, HE-congruence is the only viable form of congruence in time-independent SSEs, meaning that congruence, in general, poses no issues for inference when using the proposed parsimony refinement.

On the other hand, other congruence classes might exist in time-dependent SSEs, where different time slices of a tree have different parameters. This requires further research and is related to the concept of multiplicative closure in Markov models (41).

## Conclusion

Every SSE belongs to an infinite congruence class, making models within the class unidentifiable based on observed data alone. These congruence classes arise from the lumpability property of Markov models. For any trait-independent scenario, the congruence class includes trait-dependent models, which challenges hypothesis testing. To address this issue, we proposed a parsimony-based approach that analytically selects the simplest model within a congruence class, prioritizing the one with the fewest hidden states. Importantly, we demonstrated that the identified congruence is the only one possible in time-independent SSEs. Since our parsimony-based approach fully resolves the model selection issue, we conclude that congruence poses absolutely no challenge for inference with SSEs, unlike LP congruence in time-dependent diversification models (7).

However, model selection across congruence classes remains severely problematic, with false positive rates as high as 81-100%. The root of this error, pinpointed by congruence and lumpability, is twofold: model misspecification, to which SSEs are highly sensitive, and model proliferation, driven by the vast number of possible SSEs. Thus, our findings resolve a long-standing debate about the causes of Type 1 errors in SSEs and will be instrumental for improving inference. Due to the lack of a robust solution, we recommend exercising caution when using SSEs for hypothesis testing and following the recommendations listed above.

### Data, Materials, and Software Availability

The software, data and code necessary for reproducing the results and the figures are available on github https://github.com/sergeitarasov/Congruent-SSE-CTMC.

### HiClaSSE model

The HiClaSSE model is implemented by modifying the functions from the diversitree R package (42). Construction of congruent models was facilitated using rphenoscate R package (43).

### Congruence in MiSSE, CID4 and EHE8-C

To support the analytical findings regarding the congruence classes in MiSSE (Eq. 2), CID4 (Fig. 5B-D, H), and EHE8-C (Fig. 5H-I), we simulated 10 datasets under a CID4 model using HiSSE package (5), and performed ML inference with HiClaSSE to demonstrate that the likelihood estimates are identical. Note that the choice of the data-generating model (i.e., CID4) is irrelevant here, as congruent models remain congruent with any dataset.

### Simulations with CID4 and SEM8 models

We simulated 100 datasets under the four-parameter CID4 model (*λ*_1_ = 0.3, *λ*_2_ = 0.1, *q* = 0.2, and *µ* = 0.01; Fig. 5H) with varying numbers of tips (100, 500, 1000) using the HiSSE package (5). For each dataset, we then fitted the original CID4 model and a pool of SEM8 models constrained to only two parameters (one for speciation and state transition rates, and one for extinction). The candidate SEM8 models were selected using our optimization algorithm that traverses the SEM model space by testing 36 different rate combinations in the *Q* matrix and selecting the one with the highest likelihood. The algorithm operates by dividing the *Q* matrix into a 4×4 block matrix based on the states: *a*0*b*0, *c*0*d*0, *a*1*b*1, *c*1*d*1 (Fig. 5). For each block, it tests three distinct parameterizations, selects the best one, and then moves to the next block. We tested parameter combinations for 12 blocks, which theoretically results in a model space of 3^12^ = 531, 441 possibilities. However, the actual number of possibilities is smaller due to algorithmic inefficiencies. Despite this limitation, the approach was effective in identifying SEM8 models that fit the data better than the CID4 model. To assess model fit, we calculated Δ*IC* as *IC* = *IC*(*CID*) ≠ *IC*(*SEM*), where Δ*IC* represents either AIC or BIC (Fig. 1).

### Topological bridges

We used the data simulated in the previous step. We then fitted the original CID4 as well as COR8-NC, CID8-NC, and CID8a-NC models (Fig. 3). Finally, we calculated the expected Δ*AIC* values for each pairwise comparison.

### Empirical study

We reproduced the results of the original study (27) and fitted a set of models as shown in Table 1 using HiClaSSE. To identify the best-fitting SEM8 model, we first manually tested several candidate parameterizations of the speciation tensor and then applied the optimization algorithm described above. For fitting the EHE25-C model, we used a workaround, as HiClaSSE does not currently support models with so many hidden states. Our algorithm for constructing EHE models (SI Appendix) shows that instead of using many states, it is possible to use just a few and substitute the rest with state multipliers (i.e., *z*_*i*_). Both approaches are equivalent. First, we calculated the EHE approximation for the inferred rates in CID4 (Table 1) using the EHE algorithm (SI Appendix). Next, to simulate the behavior of EHE25-C, we used an SSE model with four states and corresponding multipliers.

We thank Matthew W. Pennell and the members of both the Tarasov and Uyeda Labs for valuable comments on this work. We credit Drägü for the phylopic shared under CC BY-SA 3.0.

ST was supported by the Research Council of Finland Grants (339576, 346294, 331631), and the three-year grant from University of Helsinki; ST partially conducted this work while a Postdoctoral Fellow at the National Institute for Mathematical and Biological Synthesis sponsored by the NSF-DBI-1300426. JCU was funded by NSF-DEB-1942717 and NSF-DBI-1661516.

## Supporting Information Text

### HiClaSSE and hidden states

In order to avoid a common misunderstanding about what hidden states are in an SSE model, we provide a theoretical background below to help understand this concept.

### The individual components of an SSE model

An SSE is a joint model that considers both observable trait evolution and diversification regimes simultaneously. To understand SSE models, including HiClaSSE, it is practical to start with the CID model, where trait and diversification are independent. The CID model is equivalent to two separate models — one for diversification and one for trait component. In other words, CID is a separable model, whose likelihood is the product of the likelihoods of the individual models. Let us consider the simplest such model, CID4, with four states, describing the evolution of a binary trait and binary diversification regime.

The trait component is a traditional CTMC, represented by a rate matrix *Q*_*t*_, with each state being observable (Eq. 1). This model is traditionally used for ancestral state reconstruction of binary traits.

The diversification component is another CTMC characterized by a rate matrix *Q*_*r*_ (Eq. 1), where the regimes are discrete states (A and B) like in the trait’s CTMC. Each state is equipped with corresponding speciation (*λ*_*A*_, *λ*_*B*_) and extinction (*µ*_*A*_, *µ*_*B*_) rates, governing the birth and death of lineages. It is convenient to think of this CTMC from a data generation perspective. When this process generates a phylogenetic tree, it transitions between regimes, and depending on the regime, a lineage can split into two lineages or go extinct according to the speciation and extinction rates. The diversification component alone is known as a trait-free MiSSE model.

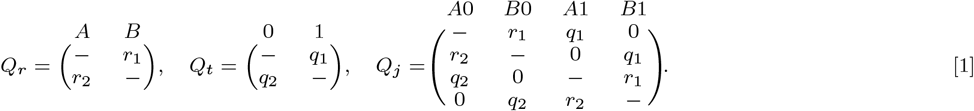

### Joint CID process

By combining the diversification and trait components, we obtain a four-state transition rate matrix *Q*_*j*_ for the joint process (Eq. 1). Each state in this matrix simultaneously belongs to a diversification regime and a trait state. Each state is associated with speciation rates (*λ*_*A*_, *λ*_*B*_, *λ*_*A*_, *λ*_*B*_) and extinction rates (*µ*_*A*_, *µ*_*B*_, *µ*_*A*_, *µ*_*B*_), following the order of states in *Q*_*j*_. As speciation and extinction rates are independent, it implies that pairs of joint states 1A and 2A, as well as 1B and 2B, have the same speciation and extinction rates as well as the *Q* matrix as in Eq. 1.

### What are the hidden states?

All states in the joint model *Q*_*j*_ are essentially hidden because we are uncertain about which states our data at the tree tips correspond to. For instance, if we know that a tip is in state 1, we do not know whether it belongs to 1*A* or 1*B*. Therefore, the observable states in our model consist of sets of hidden states. For example, an observable state 1 consists of the hidden states 1*A* and 1*B*. It is helpful to conceptualize this model as having a two-layered structure: a hidden layer that represents the actual states of the model and an observable layer to which the hidden states are mapped. The evolution takes place exclusively at the hidden layer, and its results are subsequently re-mapped to the observable layer. This two-layered approach serves as a general framework for constructing various SSE models, accommodating a wide range of scenarios. In model formalism, these two layers are modeled using ambiguous coding at the tips of tree.

For likelihood computation, the data at tips should be encoded as probability vectors, instead of integers like in character matrix. For instance, for a binary trait, data at the tips are encoded as vectors: (1, 0) or (0, 1), representing the observed states 0 and 1, respectively. In cases where the tip’s state is uncertain, we use an ambiguous coding, such as (1, 1), which implies that both states are possible.

The same approach is adopted by SSEs. For example, the CID model (Eq. 1) requires the data to be encoded as (1,1,0,0) and (0,0,1,1) corresponding to the observable states 0 and 1 respectively. For the two-state MiSSE model (e.g., *Q*_*r*_ in Eq. 1), all tips are encoded as (1, 1), indicating that all states are hidden, which emphasizes our lack of knowledge about the diversification regimes at the tips, which the model aims to infer.

### Correlated SSE models

The SSE model displaying correlated evolution between trait and diversification can be created from CID by altering rate symmetries in *Q*_*j*_ or by rewiring the association between hidden states and speciation/extinction rates. See the section “Correlated Evolution in SSE Models”.

### BiSSE model and hidden states

Any SSE model, including those without hidden states like the BiSSE model, can be viewed as an instance of the general “hidden state” framework. By modifying the mapping of speciation and extinction rates in the CID model (Eq. 1), we can create a correlated process. For example, we can set the rates as follows: speciation (*λ*_*A*_, *λ*_*A*_, *λ*_*B*_, *λ*_*B*_) and extinction (*µ*_*A*_, *µ*_*A*_, *µ*_*B*_, *µ*_*B*_) while keeping *Q*_*j*_ the same. This modified model, which we will call COR4, is congruent to the BiSSE model because the rate matrix of COR4 (i.e., *Q*_*j*_ in Eq. 1) and the diversification rates are lumpable with respect to the state partition {{*A*0, *B*0}, {*A*1, *B*1}}. So, the corresponding BiSSE model is defined by *Q*_*t*_ (Eq. 1) and the diversification rates are: (*λ*_*A*_, *λ*_*B*_), (*µ*_*A*_, *µ*_*B*_). Although the BiSSE model does not formally have hidden states, it can be viewed as a simplified version of a more complex hidden state model.

### HiClaSSE likelihood

The HiClaSSE description follows the general framework for SSE models. We will use lowercase symbols for scalars (e.g., *a*), italic-bold symbols for column vectors (e.g., ***a***), capital letters for matrices (e.g., *A*), and bold capital letters for sets of matrices, called tensors (e.g., **A**). The element-wise product, denoted as *¶*, involves multiplying each element of two arrays with the same dimensionality. The notation ***u*** *⊗* ***v***^*T*^ signifies the outer product of two vectors, where ***Xu*** represents the multiplication of matrix ***X*** by vector ***u***, and ***XY*** indicates the multiplication of matrix ***X*** by matrix ***Y***.

We consider that HiClaSSE represents a lineage-varying state-dependent birth-death process with ***k*** discrete states and the following types of events: (1) speciation events, where a lineage splits into two; (2) extinction events, resulting in the death of a lineage; and (3) state-change events, where a lineage change from one state to another.

### Speciation events

We denote by **Λ** the speciation array that represents a set of *k* matrices **Λ** = {Λ_1_, Λ_2_,…, Λ_*k*_} characterizing the rates for speciation events. Each matrix Λ_*i*_ has dimension *k* × *k*, with upper triangular entities populated with *λ* rates (others are zero). Matrix Λ_*i*_ has entities *λ*_*ijz*_, which refer to the speciation rates that occur when a lineage in state *i* speciates and produces two daughter lineages one in state *j* and another in state *z*:

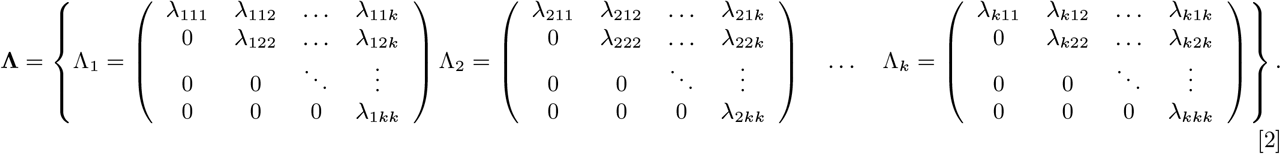

All these events fall in two categories – those simultaneously occurring with state changes and those which do not. We refer to them as coupled (e.g., *λ*_111_) and decoupled (e.g., *λ*_122_) speciation events respectively.

Note that Λ_*i*_ is intentionally an upper triangular matrix to visualize unique rate entities and avoid rate duplication (as *λ*_112_ = *λ*_121_). However, in some calculations below, we will require a symmetric speciation matrix 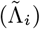 where the zeros in Λ_*i*_ are filled with rates from the upper triangle. Such matrices can be obtained through the following algebraic operation:

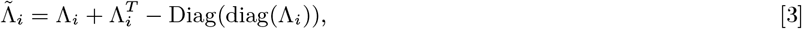

where, superscript *T* denotes the transpose of matrix, diag(Λ_*i*_) extracts the diagonal of Λ_*i*_, and Diag(diag(Λ_*i*_)) forms a diagonal matrix with the diagonal elements of A_*i*_. These symmetric matrices also form an array 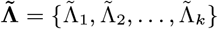

We also need to define a matrix-wise operation on the matrix arrays (Λ and 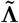). For example, a notation Λ *○****X*** indicates that each Λ_*i*_ in **Λ**should be multiplied via *○* by matrix ***X***:

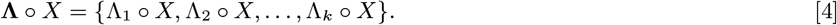

Similar matrix-wise operations apply to Λ and 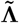 for vector multiplication (e.g., **1**^*T*^ **Λ1**). We will use the matrices ***I***_2,1_ and 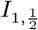 in matrix-wise operations on arrays. *I*_2,1_ is a matrix where all diagonal elements are 2s, while off-diagonals are 1s. It multiplies a diagonal element in each 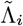by two. 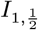 is a matrix where all diagonal elements are 1s, while off-diagonals are 1*/*2s. It divides the off-diagonal elements in each 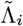 by two. These matrices are necessary to control for the number of speciation and extinction events in calculations.

### Extinction events

The extinction rates for each state are given by the column vector ***µ***:

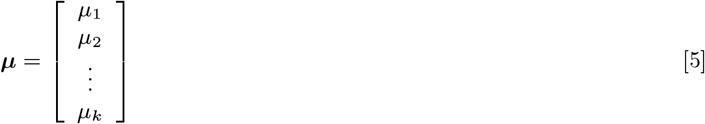

### State-change events

The instantaneous rate matrix matrix ***Q*** defines anagenetic state changes and represents a traditional CTMC. Its entities are the transition rates ***q***_*ij*_.

### Probability Terms

The probability of observing a particular branch at time ***t***, given that the branch is in state ***i***, is denoted by the scalar ***D***_*i*_(***t***). The probability of lineage extinction at time (***t***), given the lineage is in state ***i***, is the scalar ***E***_*i*_(*t*). These two entities are equivalent to ***D***_***N***,***i***_(***t***) and ***E***_***i***_(***t***) from BiSSE. Note, that we have dropped the index *N* from ***D***_***N***,***i***_(***t***) for brevity. The entities ***D***_***i***_(***t***) and ***E***_***i***_(***t***) can be presented in vector form as:

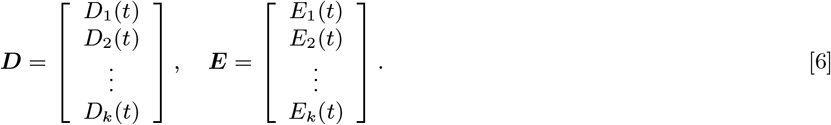

### Differential Equations and Likelihood

The probability terms evolve over time and branch according to the following system of differential equations, which is written in array notation as:

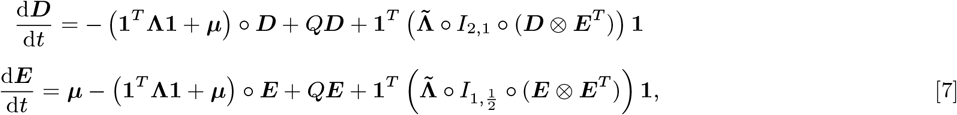

this system is equivalent to BiSSE-ness model extended for *k* states. Note, that each of the vectors, 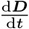 and 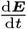, characterizes *k* differential equations.

At each ancestral node (*A*) the diversification probabilities (***D***_***A***_) calculate as:

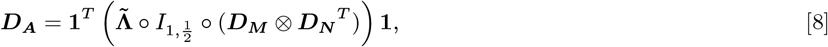

given that this node has two daughter lineages (*M* and *N*) with the the diversification probabilities ***D***_***M***_ and ***D***_***N***_ respectively. This formula is equivalent to how BiSSE-ness combines the probabilities at nodes.

The likelihood of the tree (*L*) calculates at the root as:

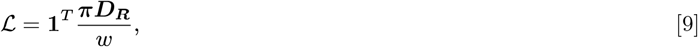

where, ***D***_***R***_ are the diversification probabilities at the root, and ***π*** is the initial vector of probabilities for each state. The term *w* (Eq. 10) conditions the overall likelihood on clade survival (1), indicating that neither of the two daughter lineages at the root is allowed to become extinct before the present. Note, both the root and likelihood equation are equivalent to BiSSE-ness model too.

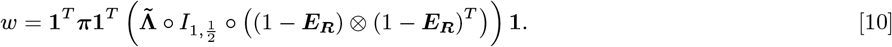

### Example: HiClaSSE2

Following Eq. (7), the HiClaSSE model with two states is defined by these ODEs:

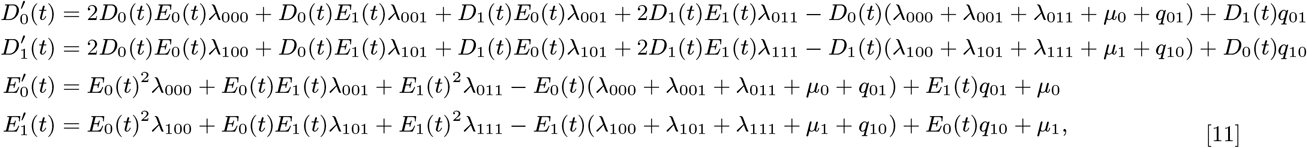

where, ***f’*** indicates 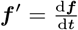

These equations (Eq. 7) are equivalent to BiSSE-ness model, and thereby ClaSSE, but have a different parametrization. The BiSSE-ness model has 10 parameters: (*λ*0, *λ*1, *µ*0, *µ*1, *q*01, *q*10, *p*0*c, p*0*a, p*1*c, p*1*a*). These parameters include two speciation rates (*λ*0, *λ*1) that are weighted by probabilities representing anagenetic or cladogenetic speciation events. The parameter *p*0*c* is the probability of cladogenetic speciation when the ancestral lineage is in state 0, which can be asymmetric (*p*0*a*) or symmetric (1 ≠ *p*0*a*). Similar logic applies to *p*1*c* and *p*1*a*. The mapping between BiSSE-ness parameters (RHS) and HiClaSSE2 (LHS) is as follows:

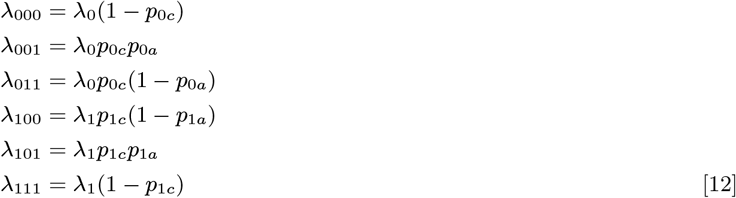

### Correlated Evolution in SSE Models

Mark Pagel proposed conditions to determine whether two binary traits should be considered correlated or independent. We can extend these conditions to SSE models. Let us take a SSE model with two speciation and extinction rates: (*λ*1, *λ*2, *λ*1, *λ*2), and (*µ*1, *µ*2, *µ*1, *µ*2), and the rate matrix:

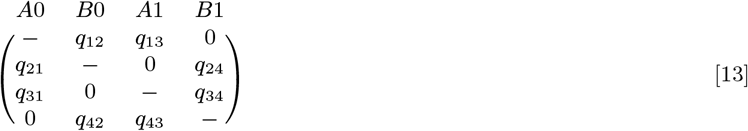

For this model to be independent (i.e., CID), the following conditions should hold: *q*13 = *q*24, *q*31 = *q*42, *q*12 = *q*34, *q*21 = *q*43. If these conditions are not met, the model implies the following dependencies:

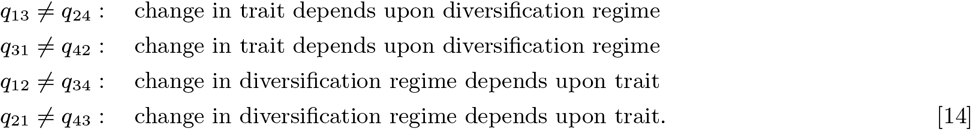

Consequently, if two different types of conditions from Eq. (14) are met simultaneously (e.g., *q*13 ≠ *q*24 and *q*12 ≠ *q*34), then the changes in both the trait and diversification regime are interdependent.

It is worth noting that there are additional conditions suggesting correlation. If any of the left diagonal rates, indicating dual transitions (e.g., *A*0 *⟶ B*1 is allowed), has a non-zero value, then a change in trait implies an immediate change in diversification regimes, and both are mutually dependent. Note that all these correlation conditions are only valid when we have at least two speciation or extinction rates (as in the CID model of this section). In the trivial case of equal speciation and extinction rates across all states, the diversification is naturally not affected by the trait since its invariable.

Non-independence can be proven by the fact that an independent SSE is separable, thereby it can be constructed from two individual models, one for trait and another for diversification. If this is not possible, then the SSE is correlated. This construction can be achieved by manually deriving the joint probabilities. Alternatively, for constructing the joint rate matrix ***Qj***, there is also a convenient algebraic operation using the Kronecker sum (⊗) that combines individual matrices.

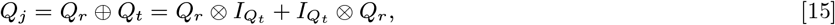

where, ***Q***_***r***_ and ***Q***_***t***_ are the individual rate matrices for diversification regimes and traits, 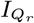 and 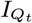 are the identity matrices for the respective *Q*’s, an*d* ⊗ denotes the Kronecker product. Consequently, all joint matrices that cannot be constructed using Eq. (15) imply correlation.

As an example, consider the models: COR8-C and COR8-NC from Eq. (**??**) (Table **??**). Their rate matrices cannot be produced by combining individual trait and diversification models using Eq. (15). Therefore, they are not separable and describe dependent evolution. COR8-NC is a distinct model, while COR8-C is congruent with CID4 (Table **??**) and shares the same likelihood.

### Lumpability Conditions for SSE models

#### Lumpability of a basic CTMC

Let us begin by discussing lumpability in the context of a basic CTMC and then extend it to SSE models. Suppose, we have an initial CTMC that describes the evolution of a four-state trait {*s*1, *s*2, *s*3, *s*4} and is defined by the rate matrix *Q*:

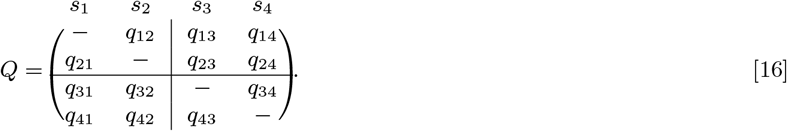

The aggregation of states at the level of the rate matrix means constructing a smaller matrix with fewer states and transitions. Suppose that we are willing to aggregate *Q* into a two-state process 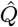, with states {*ŝ*1, *ŝ*2}, using the following partition of the initial states: *ŝ*1 = {*s*1, *s*2}, *ŝ*2 = {*s*3, *s*4}. This partition can be visualized on *Q* by splitting the rates into four partition blocks shown with a horizontal and a vertical line within the matrix in the Eq. (16). Each block includes four initial rates whose function should yield new rates in the aggregated matrix. So, the aggregated matrix should be:

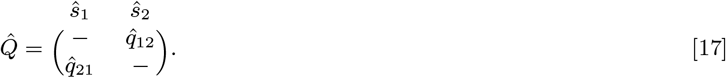

The lumpability property ensures that the transition rates of the aggregated CTMC can be calculated from the original CTMC, allowing us to model them using the aggregated process. The lumpability property described in the main text can be restated in the context of partition blocks within the rate matrix. We refer to this restatement as the “row-wise sum rule” (RWR). Both interpretations are equivalent.

The RWR implies the following: *the original CTMC is lumpable with respect to a given partition of states when the row-wise sum of rates within one partition block in the original Q is the same for all rows within the given partition block, and this property holds for all blocks in Q. The rates in the aggregated matrix represent simply the row-wise sums of the original rates*.

For *Q* in the equation Eq. (16), the RWR is maintained if the following two equalities hold:

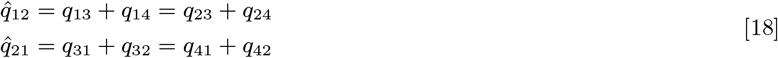

These equalities imply lumpability of *Q* for the given state partition; 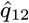 and 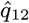 represent the new rates in the lumped process 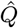, shown in the equation Eq. (17). Note, to prove lumpability, it is enough to show that the RWR holds for the off-diagonal blocks because it implies that the main diagonal blocks maintain the RWR too. This property will be helpful later, in classifying hidden expansions. In the lumped process, the initial probability vector at the tree root (***π***) should be aggregated, too: by adding up state probabilities from the original process belonging to the same partition subset.

The lumpability property is general and applies to all types of CTMCs. Lumpability does not depend on the time over which a CTMC evolves nor the initial probability vector at the tree root; if a CTMC is lumpable, it is lumpable under any value of the initial vector. If the lumpability property does not hold, then the simple form for the rate matrix for the aggregated process does not exist since it is not Markovian.

### Lumpability and backward ODEs

The previous example demonstrated lumpability for forward-in-time equations. However, in phylogenetics, likelihood is estimated using backward-in-time equations starting at *t* = 0. Let us first consider these equations for a basic CTMC from Eq. (16) to demonstrate our approach.

The system of backward-in-time equations for calculating likelihood over a tree branch can be written in matrix form as***D***^*’*^ = *Q****D***, where ***D*** is:

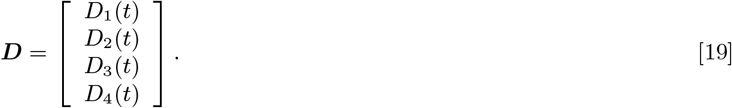

This system expands as:

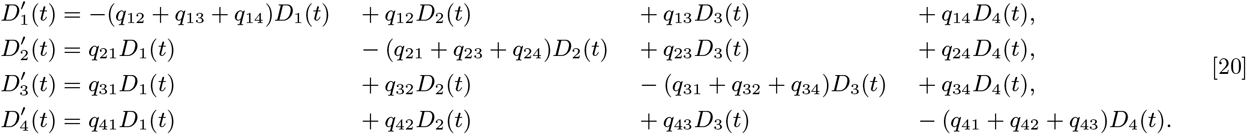

Now, we want to construct a smaller CTMC given the state partition *ŝ*1 = {*s*1, *s*2}, *ŝ*2 = {*s*3, *s*4} as above. Our approach involves first deriving a necessary condition for the smaller CTMC to be Markovian, then applying it to the system of the original ODEs, and finally deriving the relationships between the rates that are necessary and sufficient to maintain lumpability. For the smaller CTMC to be Markovian in the backward equation, it is necessary that we do not distinguish between the original states that belong to the same lumped state. Thus, the *Di*(*t*)’s belonging to the same lumped state must be equiprobable for any time *t*. In our example, this means that *D*1(*t*) = *D*2(*t*) and *D*3(*t*) = *D*4(*t*). Putting this conditions in equation Eq. (20) and after some algebra gives:

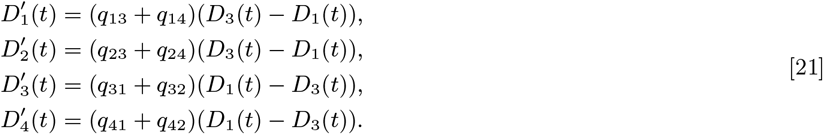

Clearly, for this system to be lumpable the same equalities as in Eq. (18) should hold. This allows us to rewrite these four equations using just two equations that characterize the lumped process from Eq. (17):

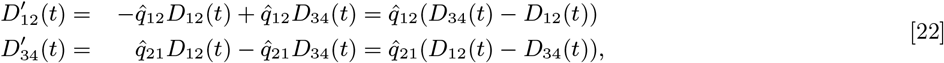

where *D*12(*t*) = *D*1(*t*) = *D*2(*t*) and *D*34(*t*) = *D*3(*t*) = *D*4(*t*); the dependencies between the rates are the same as in Eq. (18). We will employ this approach to derive lumpability conditions for SSE models.

### Lumpability Conditions for SSE

The ODEs for SSE models can become quite extensive and involve numerous parameters depending on the number of states. For simplicity, let us illustrate the method discussed in the previous section by applying it to a HiClaSSE3 with states denoted as {0, 1, 2}. Our goal is to lump these states into two states given the partition {{0}, {1, 2}}. We assume that the original SSE model has the maximum number of parameters, which include the extinction rates ***µ*** = [*µ*0, *µ*1, *µ*2], and the transition rate matrix *Q*3:

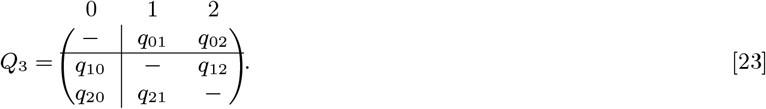

The coupled and decoupled speciation rates are specified by **Λ** tensor:

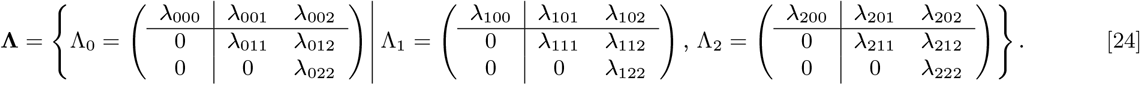

To lump the model, we set *D*1(*t*) = *D*2(*t*) and use the ODEs fromEq. (7), which yields:

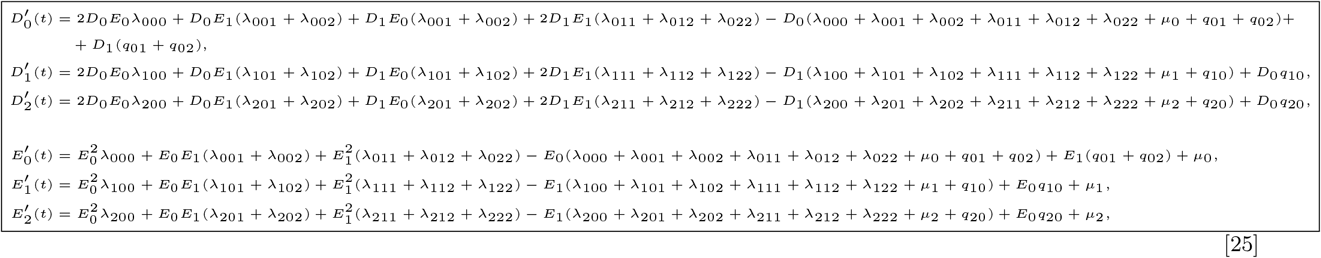

where *D*’s and *E*’s are, indeed, *D*(*t*)’s and *E*(*t*)’s, with *t* omitted for brevity. You may notice that the rates across these equations group together based on the same *D* and *E* terms. For example, for *D*^*′*^ (*t*), there is an entity *D*0*E*1(*λ*101 + *λ*102) and for *D*^*′*^ (*t*), there is *D*0*E*1(*λ*201 + *λ*202). If *λ*101 + *λ*102 = *λ*201 + *λ*202, then the the two entities are equal. Thus, we can make the HiClaSSE3 model lumpable by ensuring that all such entities are equal. According to our state partition, HiClaSSE3 is lumpable if:

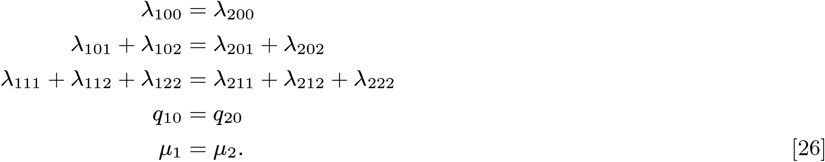

If all of these conditions are satisfied, it allows us to rewrite the HiClaSSE3 model from Eq. (25) as a HiClaSSE2 model with two states as shown in Eq. (11). The following relationships exist between the HiClaSSE3 (LHS) and HiClaSSE2 (RHS):

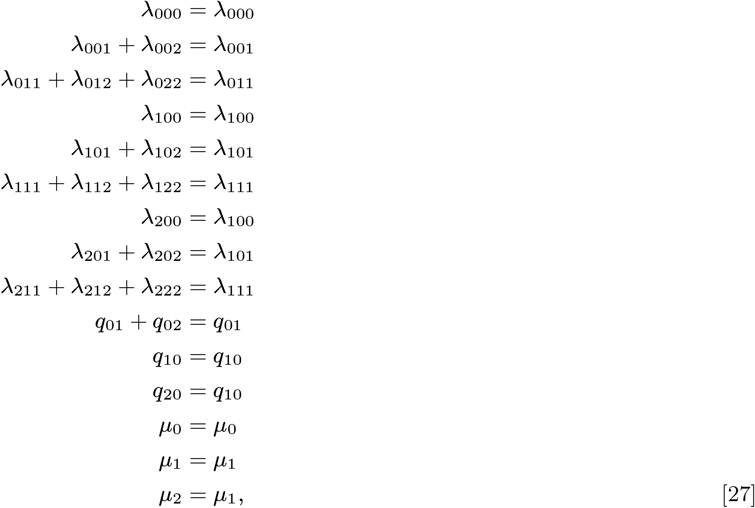

### The lumpability algorithm

For an SSE to be lumpable, each of its components: **Λ, *µ***, and *Q*, should satisfy lumpability conditions. While the lumpability conditions for ***µ*** and *Q* are straightforward and have been discussed earlier in detail, the conditions for the **Λ** tensor may be cumbersome. If we think of **Λ** not as a set of matrices but as a 3D array, then its lumpability is analogous to the RWR, but in three dimensions.

To check if **Λ** is lumpable given a certain state partition *B*, you can follow this algorithm: (1) split each A*i* matrix in **Λ** into partition blocks, as done with *Q*, according to *B* (e.g., Eq. 24); (2) Then partition **Λ** itself by grouping individual matrices into subsets according to *B* (e.g., as shown by the vertical bar after A0 in Eq. 24); (3) **Λ** is lumpable if, for each subset of matrices, the sum of rates across identical partition blocks is the same, and this condition holds for all subsets and blocks.

### Lumpability and Model Congruence

If the original model is lumpable with respect to the hidden states, then the lumped model is congruent with the original one. This can be understood by examining *D*(*t*)’s and *E*(*t*)’s. When calculating likelihood over a tree branch using Eq. (7), *D*(*t*)’s and *E*(*t*)’s belonging to the same observable state are identical due to the lumpability assumption. This implies that they are also the same at nodes and the root (Eq. 8). Therefore, lumping by aggregating hidden states essentially means that *D*(*t*)’s and *E*(*t*)’s at the root between the two models only differ in the number of states, while the likelihood values at the root (Eq. 9) are identical. Accordingly, the lumpability holds regardless of the sampling fraction, since it simply alters the relative value of *D*(*t*)’s and *E*(*t*)’s.

### Equal Rate Hidden Expansion (EHE)

The CID model used in the EHE transformation example consists of four parameters with specific parameter values. Its expansion results in EHE8-C, which has only two parameters and includes cladogenetic speciation events (Table **??**, the bottom part; and Eq. 28).

Note that in the EHE example, the rates are exact multipliers of 0.1 for demonstration purposes. This is done to simplify the visualization. If other values with many decimals were used, EHE transformations could result in rate matrices with thousands of states, making it impractical to visualize them.

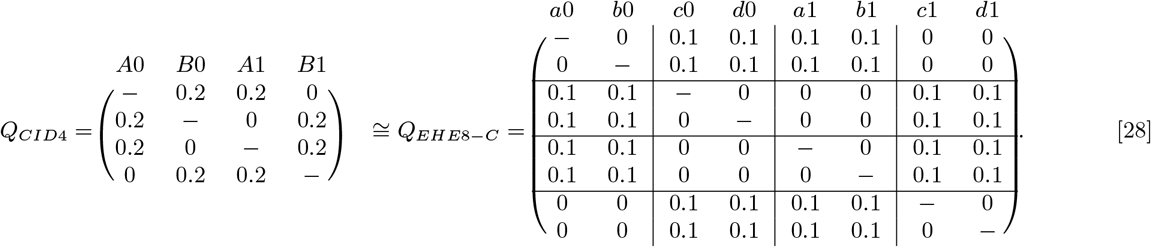

### The EHE Algorithm

The EHE algorithm expands any CID model with decoupled separation events to an EHE model. It can also be adopted for any SSE. This algorithm expands each initial state with an equal number of additional hidden states. To calculate the minimal EHE model, follow these steps:

1. Pool together the rates from *Q* (only non-negative rates) and **Λ** into a single vector. Represent them as irreducible fractions of two integers and find their least common denominator (*lcd*).
2. Define the equal rate as *r* = 1*/lcd*.
3. Find the maximum rates in **Λ**and *Q*. Denote them as *λ*_*max*_ and and *q*_*max*_ respectively.
4. Calculate the necessary number of hidden states per initial state, represented as *N*_A_ and *N*_*Q*_, corresponding to **Λ** and *Q* respectively. For the rate matrix *Q*, this number is:

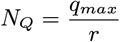

For the speciation tensor **Λ** is:

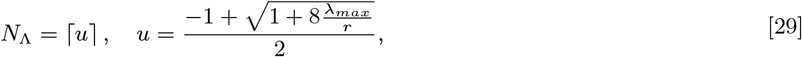

where ⌈ · ⌈ represents the ceiling function. The ceiling of a number *x* is the smallest integer that is greater than or equal to *x*. It is required since the number of hidden states is an integer. The formula from Eq. (29) derives from the number of different lambda rates 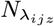 per initial CID state in the expanded model, which is:

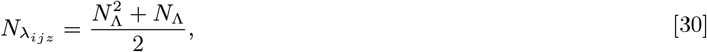

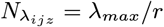setting *N*_*λ*_ = *λ*_*max*_*/r*, and solving Eq. (30) yields Eq. (29).
5. Select the maximum value to determine the final number of hidden states *N*_*h*_ needed for the observable one: *N*_*h*_ = max(*N*_*Q*_, *N*_A_).
6. Create the appropriate EHE model, corresponding to *N*_*h*_, and distribute the rates to maintain lumpability.

For the CID4 from Eq. (28) and Table **??**, the algorithm operates as follows. The three rates {0.1, 0.3, 0.2} can be represented as fractions: 1*/*10, 3*/*10, and 2*/*10; their *lcd* = 10, hence *r* = 1*/lcd* = 0.1. The maximum rates are: *qmax* = 0.2 and *λmax* = 0.3. So, *N*_*Q*_ = 0.2*/*0.1 = 2, *N*_A_ = 2, which means that *N*_*h*_ = 2, resulting in EHE having two hidden states for each initial state, as illustrated in Eq. 28 and Table **??**.

The algorithm outlined above constructs the EHE model with the minimal number of hidden states. However, if one selects a different common denominator (other than the *lcd*) and applies the same algorithm, it will generate an EHE model with a greater number of hidden states.

### The Structure of a Congruence Class

To derive parametric forms for congruent models, we classify HEs into three main types based on the number of parameters in a HE model compared to the original SSE. This classification focuses on counting parameters located in the off-diagonal blocks (referred to as off-diagonal parameters) of the *Q* matrix in the expanded model, while ignoring the total number of parameters in the model. This distinction is crucial because the parameters in the diagonal blocks are unidentifiable and can be set to any values, as they do not affect the RWR (see the section “Lumpability of a basic CTMC”). For example, setting the diagonal *q*’s in the *QCID*8 from Eq. (**??**) to 0 or any other separate parameter will not impact lumpability.

The three main types of HEs are as follows: equivalent expansion, super-expansion, and sub-expansion. These types generate models with the same, greater, and fewer off-diagonal parameters, respectively.

We also define two additional categories of HE models that are crucial for analyzing SSE dynamics. The Waiting-Time Preserving (WTP) hidden expansion does not alter the expected waiting times in *Q* between the original model and HE. These waiting times are indicated by the negative main-diagonal entries in the rate matrix. Conversely, the opposite category is a non-Waiting-Time Preserving (nWTP) expansion, which modifies the expected waiting times in *Q* between the original and HE models. For instance, *QCID*8 in Eq. (**??**) is nWTP, but if we change the *q*’s in the diagonal blocks to 0, it becomes WTP.

The parametric forms for equivalent and super-expansion models can be directly derived from *M*0 using the lumpability conditions. The total number of parameters in these models remains the same (e.g., WTP equivalent type models) or increases (e.g., nWTP equivalent or super-expansion models). Super-expansion models have the potential for an unlimited number of parameters since there is no upper bound to maintain lumpability.

On the other hand, the parametric forms for sub-expansion models depend on the specific parameter values of *M*0. This implies that each point in the parameter space of *M*0 corresponds to a unique sub-expansion model. The total number of parameters in sub-expansion models can vary: (1) it can be fewer than in *M*0 (e.g., WTP models), ranging down to two parameters (EHE), indicating that these models are nested within *M*0; or (2) it can be the same as or greater than in *M*0 (e.g., nWTP models).

